# The Effects of Acute Coordinative vs. Acute Endurance Exercise on the Cortisol Concentration

**DOI:** 10.1101/2024.03.25.586687

**Authors:** Henning Budde, Mirko Wegner, Christiane Ahrens, Bruna Velasques, Pedro Ribeiro, Sergio Machado, Thomas Gronwald, Sandra Amatriain-Fernández, Marcelle Schaffarczyk, Anett Mueller-Alcazar

**Affiliations:** Institute for Systems Medicine (ISM), MSH Medical School Hamburg University of Applied Sciences and Medical University Hamburg, Germany; Department of Health Science, Institute of Human Movement Science, University of Hamburg, Germany; Bioscience Department, School of Physical Education of the Federal University of Rio de Janeiro, Brazil; Institute of Applied Neuroscience, Rio de Janeiro, Brazil; Brain Mapping and Sensory Motor Integration, Institute of Psychiatry, Rio de Janeiro, Brazil; Panic and Respiration, Institute of Psychiatry, Federal University of Rio de Janeiro, Brazil and Laboratory of Physical Activity Neuroscience, Neurodiversity Institute, Queimados-RJ, Brazil; Institute of Sport Science, Friedrich Schiller University Jena, Germany; Institute for Cognitive and Affective Neuroscience, MSH Medical School Hamburg University of Applied Sciences and Medical University Hamburg, Germany

## Abstract

**Background:** Physical exercise interventions can cause neuroendocrine activation, which in turn increases salivary cortisol concentrations. Until now there have been no studies comparing endurance and coordinative exercise.

**Objective:** The purpose of this study was to examine the effects of these different interventions with an intraindividual comparison.

**Methods:** 61 students between 18 and 30 years of age were included and first completed a coordinative exercise and one week later an endurance exercise of the same intensity and length which was self-set on the first day, with a maximum heart rate of 64 - 76% (HRmax) over a period of 15 min. To measure changes in the hypothalamic-pituitary-adrenal axis (HPA-axis) activity, saliva samples were collected before (t1) and after exercise (t2 and t3).

**Results:** Baseline values of cortisol (t0) did not differ significantly between coordinative and endurance exercise, t(55) = .233, *p* = .816. Post hoc tests revealed that the cortisol values of the second coordinative vs. endurance time point, *t*(55) = 2.097, *p* = .040, *d* = .741, and third coordinative vs. endurance time point, *t*(55) = 3.004, *p* = .004, *d* = .735, differed significantly with large effect sizes.

**Conclusion:** The results show for the first time that coordinative exercise produced a higher cortisol release than endurance exercise of the same intensity and duration. Interventions such as coordinative exercise require higher cognitive components resulting in stronger cortisol release than endurance exercise. Thus, the type of acute exercise is a psychophysiological factor in determining the neuroendocrine stress response.

**Key points:** Cortisol is an important catabolic hormone when it comes to overtraining or burnout in an athlete. Our results show for the first time that acute coordinative exercise produces a higher cortisol release than endurance exercise of the same intensity and duration.

## 1. Introduction

Acute physical exercise influences neuroendocrine responses and can lead to an increase in cortisol concentration if the intensity of the intervention exceeds a certain threshold [1]. In addition, studies suggest that enhanced chronical physical activity (PA) facilitates neuroplasticity of certain brain structures and is associated with an improvement in neurogenesis, synaptogenesis, the release of neurotrophins and neuroendocrinological changes that are linked to benefits for cognitive and affective functions [2]. In the long term, this may be reflected in a reduction in psychosocial stress symptoms [3] and in short term lead to an improvement in cognitive performance [4]. Different modes of exercise have gained increased attention in research in recent years [5].

In the event of acute physical or psychological stress, our body’s stress response essentially occurs via two axes: the sympatho-adrenomedullary system puts the body into an alarm state when it is exposed to stress and ensures an increase in heart and respiratory rate by releasing adrenaline and noradrenaline [6]. The hypothalamic-pituitary-adrenal axis (HPA-axis) forms the second stress axis, which is primarily characterized by the release of the stress hormone cortisol [7]. Cortisol has multiple effects on metabolism [8], the immune system and inflammatory processes [9], and is the most important hormone of the HPA-axis alongside corticotropin-releasing hormone (CRH) and adrenocorticotropic hormone (ACTH). Acute physical exercise is acknowledged as an immediate stressor for the body and affects the secretion processes of several endocrine tissues and the subsequent release of hormones like cortisol [10].

The circadian rhythm of cortisol is stable and is only slowly influenced by external conditions. During acute physical stress, a peak secretion of cortisol can be observed approximately 20 to 30 minutes after the onset of the stimulus [11]. If the organism is unable to terminate the stress response, e.g. in the case of chronic stress exposure, stress-adaptive mechanisms can lead to pathological changes. The role of the endocrine stress system can be seen in the pathophysiology of various mental illnesses such as depression [3], burnout [12], or PTSD [13]. Also physical exercise can induce changes on the functioning of the HPA-axis, provided that a certain intensity threshold is exceeded [3, 14].

Physical exercise refers to targeted, planned and structured forms of PA [15]. Physical exercise should be differentiated into acute physical exercise (single bout) and chronic physical exercise (repeated bouts of physical exercise or exercise training). The intensity and duration of the activity is considered the primary physiological factor for determining the neuroendocrine stress response [10]. Adult subjects showed a linear increase in salivary cortisol concentration over a period of 15 minutes at a maximum exercise intensity of 70-85% maximum Heart rate (HRmax), whereas no significant changes were observed at lower exercise intensities [14]. In connection with psychosocial stress, the cortisol response appears to be even more pronounced [16]. It can therefore be assumed that, in addition to the intensity of the stressors, the type of stressor itself also has an influence on cortisol levels and that psychosocial stress can increase cortisol levels significantly more than purely physical stress of moderate intensity (65-75 % HRmax) [17].

Salivary cortisol responses show great intra- and inter-individual variability and are determined by various factors [18]. Personality traits, age, gender, endogenous and exogenous sex steroid levels, genetic predispositions or critical life events and in particular negative childhood experiences are worth mentioning in this context [19]. Several studies have shown that chronic endurance training (which is defined as planned, structured, repetitive, and purposeful PA leading to a change in fitness [20]). can effect cortisol concentration in the long term [21] and thus has both a preventive and therapeutic effect in reducing the experience of stress [22]. This shows that high levels of PA are related to a lower physiological stress reactivity. The immediate neuroendocrine stress responses induced by exercise are not yet fully understood. Results of studies comparing different types of *chronic* interventions showed that coordinative exercise led to a greater reduction in cortisol concentrations and stress levels compared to chronic endurance exercise [21]. However, the immediate stress responses produced by *acute* exercise interventions are still not clear specially regarding how the acute responses differ depending on the type of exercise performed.

Based on current research, the question of whether there are direct effects of different acute interventions of the same intensity (i.e. with the same cardiovascular load) on the cortisol concentration will therefore be investigated. A distinction is made between acute physical exercise of a coordinative nature and acute physical endurance exercise. As higher cognitive abilities will be required during the coordinative exercise (e.g. the ability to react and concentrate)[23], both physiological and psychological stressors are at work here. For this reason, it is assumed that stress load will be higher. Thus, the type of acute physical stress could be a physiological factor in determining the neuroendocrine stress response. To our knowledge, there have been no studies on whether cortisol concentration differ between an acute coordinative exercise and an acute endurance exercise of the same intensity and duration. The reaction to physical interventions is demonstrably individual and depends on many influencing factors [24]. For this reason, the intra- and inter-individual variability should be given special consideration here [5] and influencing factors such as gender, PA and will be investigated exploratively.

## 2. Material and Methods

### 2.1 Participants

The G*Power analysis (α = 5%, 1-β= 95%, d = 0.95) resulted in a required sample size of 60. The participants were informed on the characteristics of their participation, agreed to participate and signed the informed consent. The study was approved by the ethics committee of the Medical School Hamburg (MSH-2021/131) and is conducted in accordance with the latest revision of the Declaration of Helsinki (2013). Further inclusion criteria for participation in the study were the absence of mental or physical impairment, drug or alcohol dependence, acute psychotic illness and a BMI of < 30 (overweight). Based on these criteria, no participant had to be excluded. Both of the exercise sessions had to be accomplished with a moderate intensity between 64 – 76% HRmax [25]. In addition, we asked the participants about their perceived social support and their satisfaction with their everyday social relationships and measured their self-perceived stress. Motor fitness and PA were also assessed. A total of 61 people between the ages of 18 and 29 took part in the study. 32 were female. The average age was 21.8 years (SD = 3.09, N = 61).

### 2.2 Study design

For the present study, a moderate exercise intensity just before the vigorous range was aimed for, based on Garber’s classification of exercise intensity [25]. The intensity of the different exercise sessions should be between 64 - 76% HRmax. According to Tanaka’s formula: HRmax = 208 - 0.7 x age [26], this corresponds to approximately 140 to 145 beats per minute (bpm). Additionally we assessed the Rate of Perceived Exertion (RPE) using the Borg Scale, a numerical scale between 6 and 20 to record the subjective feeling of exertion [27], which should correspond to a score between 13 and 14 (somewhat strenuous to strenuous). The HR (bpm) was measured before and during each intervention. The standardized individual target HR and the RPE of the coordinative intervention (Co) determined the intensity of the endurance intervention (En) and were continuously recorded and monitored with a HR monitor (Polar M430, Kempele, Finland).

After registering, the participants were instructed not to exercise 24 hours and not to eat for two hours before the start of the exercises. Each participant took part in two interventions, which took place between 2 p.m. and 4 p.m. seven days apart. In the first session, questionnaires were used to collect demographic data PA as well as the current state of loneliness which took about 20 minutes. During this time, participants were able to calm down. A coordination exercise (Co) lasting 15 minutes was initially performed. Using a coordination ladder 10 cm above the floor, the subjects were asked to complete various step and jump sequences After a washout phase of seven days, the second intervention was carried out at the same time in form of endurance exercise (En) with the same intensity for the cardiovascular system. Again, the subjects first completed a questionnaire in order to acclimatize for about 20 minutes. After putting on the HR measuring devices, the subjects had to run for 15 minutes, while getting acoustic feedback about the HR level they should achieve (second session). The rate the RPE was again assessed using the Borg scale [27]. The following questionnaires were used:

A demographic questionnaire contained physiological parameters (height, age and weight) as well as questions on the subject’s personal circumstances.

The Godin Leisure-Time Exercise Questionnaire (GLTQ) [28] was used to assess PA. The GLTQ determines the level of PA and has been shown to correlate with the level of physical fitness [29].

FFB-mot [30] was used to assess motor fitness. The subscales of the FFB-mot are the components strength, endurance, flexibility and coordination.

In addition, before the second intervention (En), the subjects’ subjective experience of loneliness was assessed using the UCLA Loneliness Scale [31].

The planned study has a quasi-experimental, „pre-post-intervention” design with repeated measurements [32]. The intraindividual variability of the cortisol concentration describes the different responses of the same subject to two different interventions at three different measurement times. The timing of both interventions was exactly the same. In order to standardize the intensity of the endurance exercise the coordinative exercise had to be performed first. A randomization of the subjects was therefore not possible. See Figure 1.

**Fig 1.**
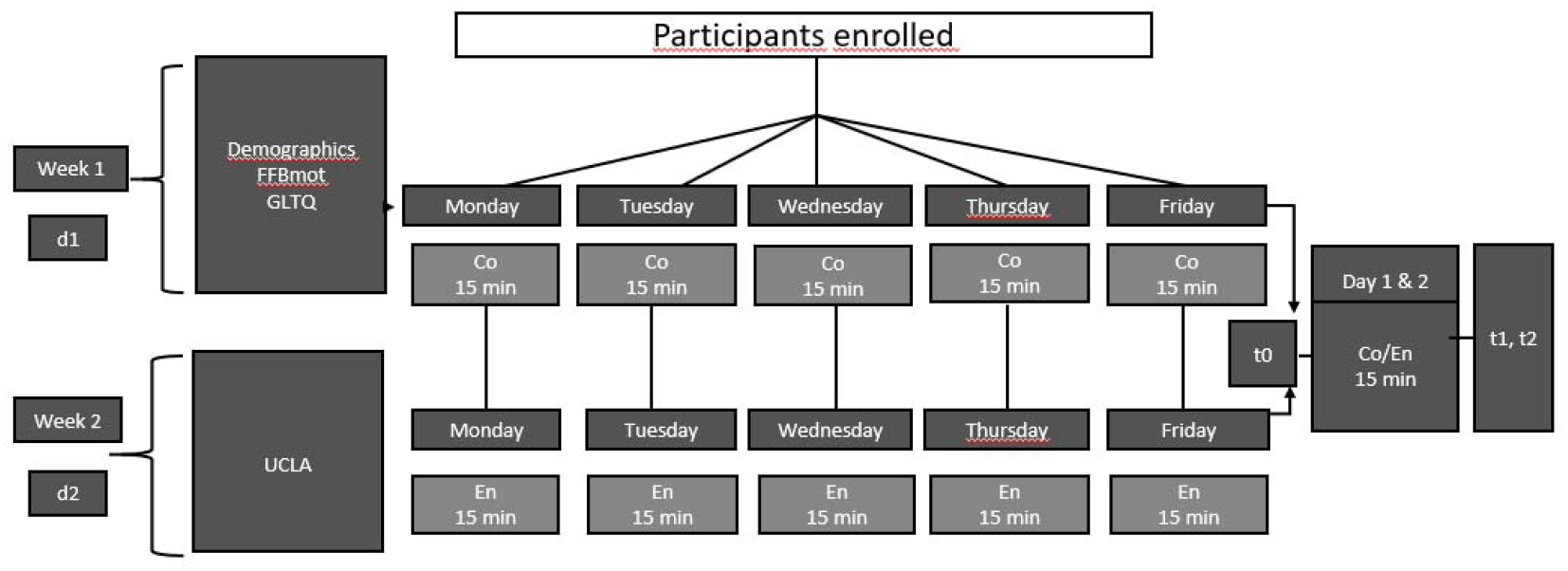
Explanation of the study design (d1/d2 = day of intervention, Co = coordinative exercise, En = endurance exercise, t0,1,2 = time of measurement, Godin Leisure-Time Exercise Questionnaire (GLTQ) [28], FFB-mot [30], UCLA Loneliness Scale [31])

The first sample was taken immediately before the intervention (t0), the second sample five minutes after completion (t1) and the third sample 30 minutes after the end of the exercise (t2). Saliva samples were collected using the SaliCap system (IBL Hamburg) and stored at minus 20°C until laboratory analysis. The saliva analysis method is considered a reliable non-invasive way of determining stress-related adrenal hormone levels [33]. The concentration of cortisol in the saliva correlates closely with the concentration of free cortisol in the blood, and it has been confirmed that the measurement in the saliva is a reliable tool for investigations of HPA activities [34]. The biochemical analysis was carried out at the University of Technology Dresden using immunoassay analysis. Intra- and inter-assay coefficients of variation were 1.8 and 2.9 %, respectively.

### 2.3 Data Analysis

Due to non-evaluable cortisol samples, two participants had to be excluded. Three others did not complete the exercise with the required intensity and were also excluded. The sample for the calculation comprises 56 participants (25 male and 31 female).

For two of the six measurement points a total of 5 samples were missing. Overall this was 1.5% missing data which allowed for imputation techniques. Multiple imputation techniques were employed using SPSS to complete the data set using the mean of five different random models. As a result complete cortisol values for all six measurement points and all 56 participants could be analyzed. For further analysis the cortisol values were log-transformed because they were not normally distributed.

The main analysis of the changes in cortisol release after both interventions (acute coordination and endurance exercise) was carried out using an ANOVA with repeated measures. This was carried out here as a special form of analysis of variance due to the presence of linked measurements or data with repeated measurements. Data were tested for sphericity and if present corrected is first tested using the Mauchly test. Effect sizes were calculated in the ANOVAS, significant results are reported by Eta-square (η^2^).

In addition, the influence of the variables gender, motor fitness (surveyed by questionnaire FFBmot) and PA (questionnaire GLTQ) on the differences between the individual measurements was exploratively examined and considered as covariates in the context of the ANOVA. The influence of gender, *F* (1,50) = 0.959, *p =* .*332*, η^2^ = 0.019, motor fitness (FFBmot), *F* (1,50) = 1.91, *p = 0*.*172*, η^2^ = 0.037, and PA *F* (1,50) = 0.029, *p =* .*866*, η^2^ = 0.001, were not significant. However, the measured cortisol concentrations were not influenced by any of these variables. The subjective feeling of loneliness (questionnaire UCLA) (F (1,50) = 0.022, *p* = .883, η^2^ = 0.001) also showed no influence on the measurement results. Therefore, in the following, analyses for Cortisol were reported and analysed without any covariate.

T-tests (two-sided) for dependent samples were calculated to test whether the cortisol concentration differed between the two interventions (Co/En). The prerequisite for the application of this method is normal distribution of the samples. The effect sizes are reported as Cohen’s d. In some cases, more than 10% of the data points for the variables under consideration are below the 5th percentile or above the 95th percentile.

## 3. Results

Preliminary analyses suggest that none of the demographic variables (age, gender, BMI, physical activity, mood) affected the results significantly and were thus not included as covariates in the ANOVA. Furthermore, baseline values of cortisol (t0) did not differ significantly between coordinative and endurance exercise, t (55) = .233, *p* = .816.

Mean values and standard deviations are shown in Table 1. Repeated measure ANOVA revealed a main effect of the measurement point, *F* (2) = 22.472, *p <* .*001*, η^2^ = .290, and the type of exercise, *F* (1) = 5.587, *p =* .022, η^2^ = .092. The interaction of measurement points and type of exercise was also significant, *F* (2, 110) = 4.322, *p =* .016, η^2^ = .073 (see Figure 2).

**Table 1.** Means (M) and standard deviations (SD) for cortisol (nmol/l) at pre- and post-test I and II for Co exercise as well as En exercise (n = 56)

| Measurement | Co |  | En |  |
| --- | --- | --- | --- | --- |
|  | *M* | *SD* | *M* | *SD* |
| *Cortisol (nmol/l)* |  |  |  |  |
| pre (t0) | 3.52 | 2.69 | 3.24 | 1.98 |
| post I (t1) | 3.24 | 2.42 | 2.54 | 1.45 |
| post II (t2) | 5.41 | 5.73 | 3.70 | 2.53 |

**Fig 2.**
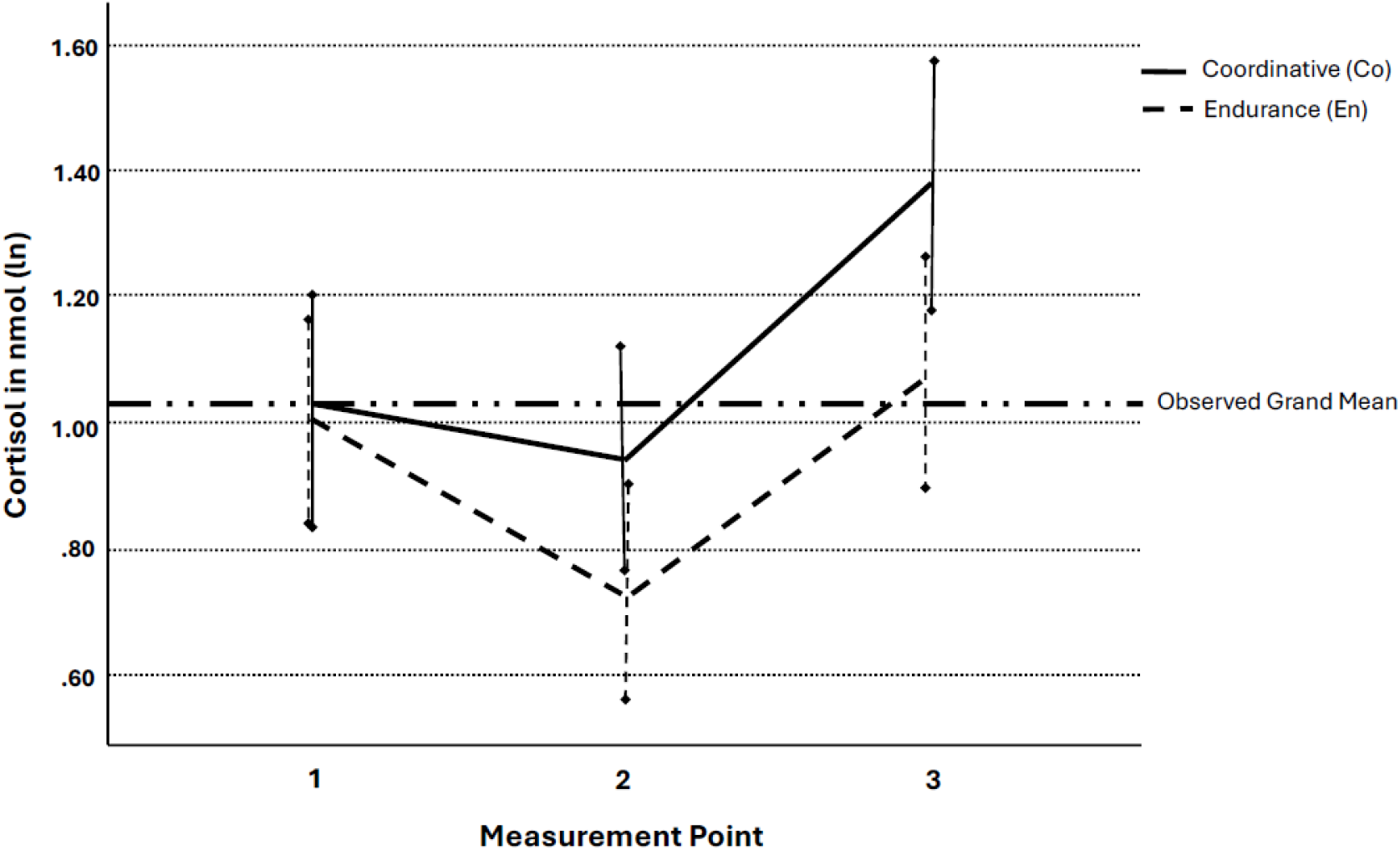
Changes in cortisol concentration after the interventions coordinative (Co) compared to endurance (En)

Post hoc tests revealed that the cortisol values of the second coordinative vs. endurance time point, *t* (55) = 2.097, *p* = .040, *d* = .741, and third coordinative vs. endurance time point, *t* (55) = 3.004, *p* = .004, *d* = .735, differed significantly with large effect sizes. Moreover, within the coordinative exercise condition, cortisol levels at time point three were significantly higher than in time point one, *t* (55) = −3.858, *p* < .001, *d* = .684, and two, *t* (55) = −6.814, *p* < .001, *d* = .484. In the coordinative condition, cortisol levels at time point one and two did not differ significantly.

In the endurance exercise condition, cortisol levels significantly differed between time point one and two, *t* (55) = 4.000, *p* < .001, *d* = .519, and between time point two and three, *t* (55) = −5.631, *p* < .001, *d* = .469. Cortisol levels did not differ significantly between time points one and three. The within conditions showed medium effect sizes.

During the coordinative exercise session, participant’s mean HR was not significantly different from the predefined HR target (144 bpm; 142 bpm). The mean HR during the endurance exercise was 143 bpm.

## 4. Discussion

Our main result was already hypothesized before the study. We saw that more cortisol is released in the aftermath of acute coordinative exercise of moderate intensity than due to an acute endurance exercise of the same intensity and duration. The endurance intervention was carried out at the same time seven days after the coordinative exercise, between 64 and 76% HRmax [27] lasting 15 minutes.

We explain our acute results with the effect we observed when comparing different types of *chronic* interventions (Cortisol Awakening Response before and after a training 3 times a week over 10 weeks). Showing that coordinative training led to a greater reduction in cortisol concentration compared to an endurance training [21]. According to Athanasiou et al. [35] regular acute exercise leads to adaptations that are reflected in a reduced physiological stress response. It would therefore be expected that enhanced cortisol concentration acutely results in a reduced concentration after a chronic treatment. If this homeostatic threat appears chronical it might lead to a lower response compared to the pretraining levels. Rehfeld et al. [36] observed a significant elevation in both white and gray matter, mainly in the cerebellum after a coordinative compared to a noncoordinative training.

It is possible that different neurobiological signaling pathways are activated by different exercise interventions [21]. The coordinative intervention places more demanding cognitive requirements on movement accuracy than endurance exercise [23]. Budde et al. [23] observed significantly higher attention after 10 min of coordinatively demanding exercise compared to endurance exercise. Coordinative exercise requires perceptual and higher-level cognitive processes, such as attention, that are essential for mapping sensation to action and ensuring anticipatory and adaptive aspects of coordination [37]. This may show that coordinative interventions lead to a pre-activation of cognitive neuronal networks [23], which is reflected in endocrine responses.

After the interventions, cortisol concentration was significantly increased in the third measurement (t3) only after the Co condition, which was performed 30 min after the end of the intervention, compared to t1. This result is not completely in line with those of comparable studies. After acute physical running exercise of a higher intensity (70-85% HRmax) over a period of 12 min in younger students, an increase in cortisol concentration could be observed in between 5 minutes after the exercise [1] whereby the intensity and duration of the exercise play a significant role in the cortisol secretion, thus cortisol was not significantly elevated after 5 min post 15 min of acute exercise in 14 year old’s (65-75% HR max) [17]. According to many studies age is an important factor in this context. In adults cortisol is first secreted a few minutes after the stressor before it then rises continuously until the peak of secretion is reached after approximately 20 to 30 minutes after the session of the exercise [11]. We also saw this rise, however, we could not show a significant differences between young and old.

In addition to the environmental influencing factors already listed, particularly pronounced personality traits such as implicit affiliation motive or power motive [19, 38] could also be attributed to affect the cortisol response.

With regard to gender [18], there was no difference to be found in cortisol secretion.

Due to our research question, we were unable to use an randomized controlled trial design [39]. We first had to determine the HR during the coordinative exercise in order to determine the intensity of the second exercise. This is more difficult with a coordinative exercise than with an endurance exercise. It therefore needed to be determined first. The lack of random assignment is the major weakness of the quasi-experimental study design [40]. However, this was unavoidable. The participants were exercising in an interindividual comparable intensity according to Garber et al. [25], we calculated their HRmax using a formula [26] and did not explicitly test their HRmax for example, with the shuttle run test [41]. However, the used questionnaire generates a reliable measurement of the motor fitness including the subscales cardiorespiratory fitness/endurance and gross motor coordination [30]. The PA-questionnaire (GLTQ) [28] we used is significantly associated with fitness of the participants [29].

## 5. Conclusion

The novelty of this intraindividual study is that it shows for the first time that after an acute exercise intervention of moderate intensity (64-76% HRmax over a period of 15 min), cortisol levels increased more under the coordinative conditions than under the endurance conditions. These results indicate that different modes of acute exercise cause different psychophysiological HPA activities, which could be of interest for future stress research and might be extrapolated to other clinical areas focused on stress responses (like exercising with a motor handicap).

## Statements and Declarations

### Data availability statement

The data that support the findings of this study are openly available in [repository name] at [URL], reference number [reference number].

### Availability of Data and Materials

Not applicable.

### Contributions

The authors HB, BV, PR, SM, TG, SA-F, MS, AM-A contributed to the design and implementation of the research, HB and CA conducted the experiment. HB, MW, CA did the analysis of the results and all were writing the draft of the manuscript. Again all authors read and approved the final manuscript.

### Corresponding author

Correspondence to Henning Budde.

### Ethics declarations

#### Ethics Approval and Consent to Participate

The participants were informed on the characteristics of their participation, agreed to participate and signed the informed consent. The study was approved by the ethics committee of the Medical School Hamburg (MSH-2021/131) and is conducted in accordance with the latest revision of the Declaration of Helsinki (2013).

## Acknowledgement

The draft is published on a preprint server: Budde, H., Wegner, M., Ahrens, C., et al. The Effects of Acute Coordinative vs. Acute Endurance Exercise on the Cortisol Concentration. Preprint at doi: https://doi.org/10.1101/2024.03.25.586687 (2024).

